# *dmrad51/spnA* mutant exhibit defects during somatic stages of developmental and show enhanced genomic damage, cell death and low temperature sensitivity

**DOI:** 10.1101/180737

**Authors:** Chaitali Khan, Sonia Muliyil, Champakali Ayyub, B J Rao

**Affiliations:** Department of Biological Sciences, Tata Institute of Fundamental Research, Colaba, Mumbai 400005, India; Current affiliation: Sir William Dunn School of Pathology, University of Oxford, South Parks Road, Oxford OX1 3RE, UK

## Abstract

<u>H</u>omologous <u>R</u>ecombination (HR) is one of the key pathways to repair <u>D</u>ouble <u>S</u>trand <u>B</u>reaks (DSBs). Rad51 serves an important function of catalysing strand exchange between two homologous chromosomes in the HR pathway. In higher organisms, Rad51 function is indispensable with its absence leading to early embryonic lethality, thus precluding any mechanistic probing of the system. In contrast, absence of *Drosophila* rad51 (Dmrad51/spnA) has been associated with defects in female germline causing ventralization of the egg, without any reported detrimental consequences to *Drosophila* somatic tissues. In this study, we have performed a systematic analysis of somatic development of *dmrad51* null mutant flies by using genetic complementation between multiple *dmrad51* alleles. Our current study, for the first time, uncovers the requirement of Dmrad51 in somatic tissue maintenance at both larval and pupal stages. Also, we show that *dmrad51* mutant exhibit patterning defects in abdominal cuticle in the stripes and bristles, while there appears to be only subtle defects in the adult wing and eye. Interestingly, *dmrad51* null mutant and other alleles show discernible phenotype of low temperature sensitivity, suggesting a role for Dmrad51 in temperature sensitive cellular processes, which thus presents an elegant system for probing temperature sensitive cellular/tissue responses that ensue when a mutation leads to the loss of protein expression (null mutant) rather than its altered protein structure. In summary, our study describes the role of Dmrad51 during somatic stages of development and provides a viable model system to study Rad51 function in a cellular process.

## INTRODUCTION

Our genome faces constant assaults owing to exogenous damaging agents as well as endogenous damages from DNA replication and transcription stress (Helleday et al. 2014). Of all types of DNA damages, DSBs (<u>D</u>ouble <u>S</u>trand <u>B</u>reaks) are the most deleterious ones and require complex mechanisms of repair (Jeggo & Lo 2007). In a cell, DSBs are repaired by two major pathways: Non Homologous End Joining (NHEJ) and Homologous Recombination (HR). HR is known to be relatively more precise but slow, largely restricted to S-G2 phase of the cell cycle, where the repair of a DSB requires a copy of the sister chromatid as a homologous template. In contrast, NHEJ occurs throughout the cell cycle, and is relatively faster, but error prone (Mao et al. 2017; Mao et al. 2008). Key steps in Homologous Recombination include homology pairing and strand invasion, during which the broken strand utilizes the information on its intact sister chromatid to repair the break. Rad51 is one of the highly conserved key players of HR, belonging to a family of DNA binding proteins with ATPase activity, which is required for the formation of active nucleofilament on ssDNA that mediates strand invasion on homologous duplex DNA (Petrini et al. 1997; Baumann & West 1998; Kelso et al. 2017). In yeast, the absence of rad51 is known to cause defects in cell cycle checkpoints and meiosis, but is not essential for cellular viability (Shinohara 1992). However, interestingly, Rad51 function becomes critical for survival in higher organisms. For instance, *rad51-/-* knockout mice exhibit early zygotic lethality at 1-2 cell stage (Shinohara 1992; Tsuzuki et al. 1996; Translocations et al. 2001), where requirement of Rad51 for cellular viability precluded its further analysis during the later stages of development.

*Drosophila* rad51 (Dmrad51 known as *spnA*) along with other DNA repair proteins was identified in a screen for maternal effect genes involved in egg pattering, giving mutants the name spindle like (spn), since eggs from these flies showed a spindle like morphology (González-Reyes et al. 1997; Ghabrial et al. 1998). It was interesting to see how the absence of repair proteins led to polarity defects in germline, thereby, indicating, for the first time, a potential crosstalk between cellular signalling and DNA repair pathways. Further, it was demonstrated that in the germline of these mutants, meiotic breaks caused by *mei-W68* endonuclease fail to repair, leading to persistence of activated DNA damage checkpoint (ATR/*mei-41*) and Chk1 (*grape*). Constitutive activation of DDR signalling in turn leads to miss-localization of EGFR ligand “gurken”, thus resulting in ventralization of the egg (Abdu et al. 2002; McCaffrey et al. 2006). Dmrad51 is also known to be expressed in somatic tissues, where its absence does not seem to affect survival rates, but has been shown to cause higher IR sensitivity (Staeva-Vieira et al. 2003; Yoo & McKee 2005). Presence of viable mutant of *Dmrad51* provided an opportunity to investigate the role of rad51 in multicellular organism; however, so far not much is known about how Dmrad51 is involved in somatic development and tissue maintenance.

In the current study, we analysed *dmrad51* mutant, for defects in somatic tissue development, by performing genetic complementation between known alleles of *dmrad51*, and corroborated the same with Dmrad51 RNAi. Our study brings out distinct patterning defects in the somatic tissue of *dmrad51* mutant; especially, in the adult abdominal cuticle, manifested as abnormal stripes along with defective bristle patterns, thus opening up interesting questions on the possible role of DNA repair components in tissue pattering and signalling. However, there were only subtle defects in the adult eye and wing of the *dmrad51* mutant. Interestingly, the larval primordial structures (wing and eye imaginal discs) of these mutant flies seem to incur significant levels of DNA damages followed by cell death. All these collectively suggest an important possibility for the role of Dmrad51 in somatic tissues. The absence of phenotypic defects in adult wing and eye, hints towards a possibility of robust compensatory mechanisms in these tissues. Interestingly, our recent study uncovered an unusual upstream regulation by initiator caspase, Dronc via its non-canonical function, in promoting ATM-mediated DDR function that might also contribute to near-normal phenotype in *dmrad51* wing discs (Khan et al. 2017).

Intriguingly, we also observed an unusual sensitivity of *dmrad51* mutant to low (18°C) temperature, with an exposure to high (29°C) temperature leading to a rescue of the phenotype, even amongst the protein null mutant. Low temperature sensitivity in the absence of protein product renders classical cold sensitive (-cs) mutants very distinct from protein structure altering cold-sensitive mutants. In summary, our study offers a characterization of *dmrad51* mutant for future mechanistic investigations involving tissue homeostasis and DNA repair along with the cellular functions of Rad51, using a rich repertoire of genetic tool-kit in *Drosophila*.

## RESULTS

### Null allele of *dmrad51* (*spnA093^A^/093^A^*) leads to an enhanced lethality during somatic stages of development in *Drosophila*

Previous studies have implicated Dmrad51 to be important for meiotic processes in the *Drosophila* germline where its absence lead to defects in egg patterning and eventually to organismal infertility (Staeva-Vieira et al. 2003). Also, it was known that Dmrad51 was not essential for organismal viability but instead that its absence was associated with only an enhanced sensitivity to ionizing radiation and genotoxic stress during larval stages (Staeva-vieira et al. 2003; Yoo 2006). However, so far, there have been no reports ascribing any phenotypic defects to *dmrad51* mutant in somatic tissues during development. In this study, by using different alleles of *dmrad51* [*spnA093^A^-null allele, spnA003-hypomorphic allele, spnA050-uncharacterized allele* and a deficiency line (99D1-99D2; 99E1)] (Tearle & Nusslein-Volhard 1987; Staeva-Vieira et al. 2003), we investigated in detail the requirement of Dmrad51 in development during somatic stages (Fig. 1A and B). Mutant alleles used in the current study were validated by genomic sequencing (Fig. 1C). The results obtained from these mutant were further confirmed using tissue specific RNAi knockdowns.

**Fig. 1.**
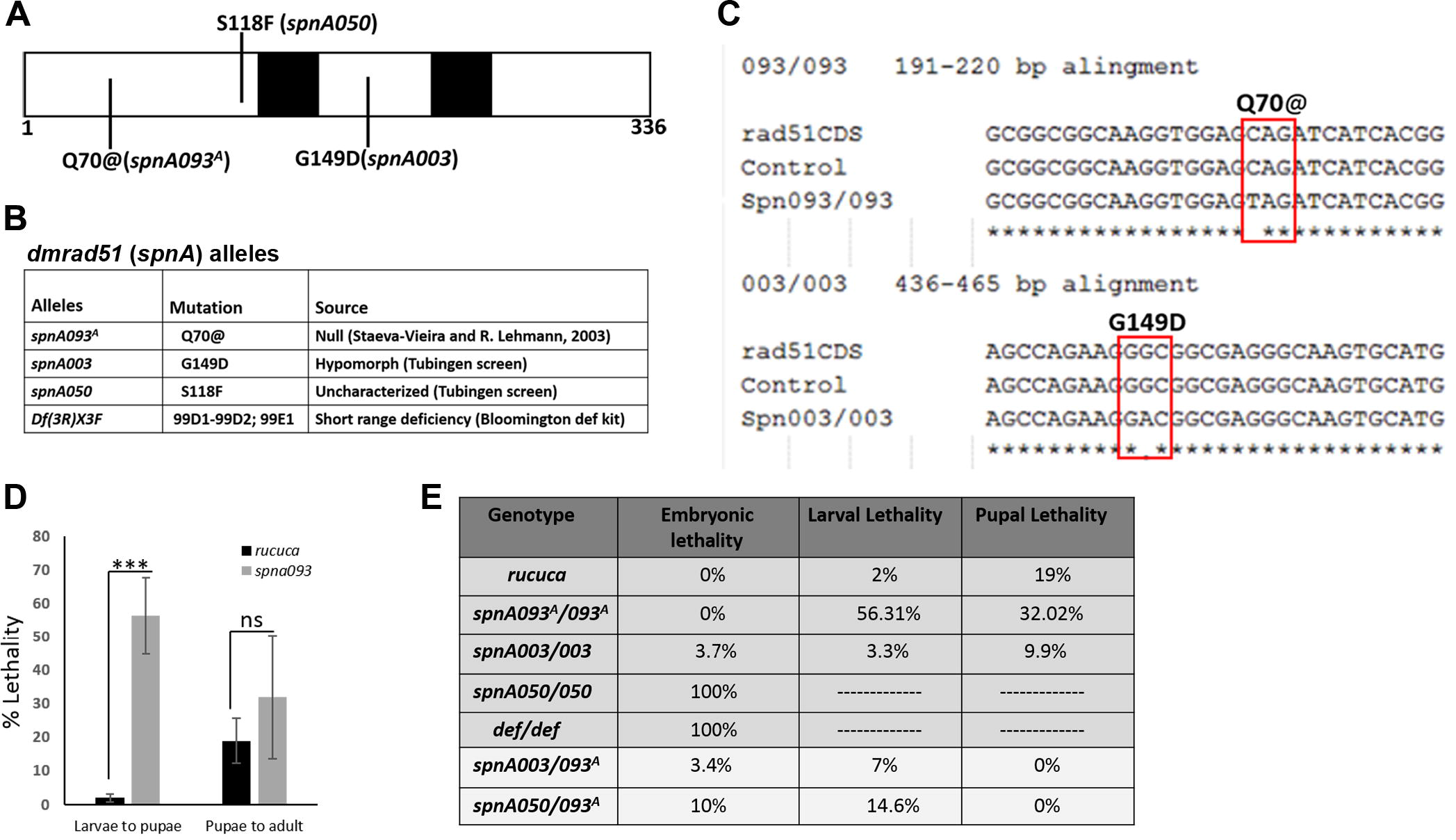
Null allele of *dmrad51 (spnA093^A^/093^A^*) leads to enhanced lethality during somatic development. (A) Dmrad51 Walker motif domain (black box) and the sites of mutation marked (adapted from Staeve-Viera et al. 2003). (B) A table of different alleles with their respective mutations and their sources. (C) Sequence validation of alleles of *dmrad51* by genomic PCR. (D and E) Developmental lethality (Expressed as percentage of total flies) associated with null mutant (*spnA093^A^*) during larval and pupal stages (D). Table shows extent of lethality for different alleles of *Dmrad51* and for trans-heterozygote with *spnA093^A^* (E).

We show that the absence of Dmrad51 (*dmrad51* null allele-spnA093^A^) causes significant lethality during larval and pupal stages in comparison to the isogenic control (*rucuca*) (Fig. 1D and E). In contrast, hypomorphic alleles (*spnA003*) did not show any lethality, while (*spnA050*) was 100% embryonic lethal and, as expected, *def/def* line showed complete embryonic lethality (Fig. 1E). Further, we performed genetic complementation analysis using the *dmrad51* alleles for lethality by using trans-allelic combinations with *spnA093^A^*, where we observed that lethality was reduced across various combinations, perhaps owing to the presence of partially functional forms of Dmrad51 (Fig. 1E). All these results put together, led us to conclude that the absence of Dmrad51 confers partial lethality during somatic stages of development, suggesting an important role played by Dmrad51 in somatic tissue maintenance, in addition to its well-known role in female meiosis, during the repair of endogenous DNA damages.

### *dmrad51* mutant show defects in adult abdominal cuticle patterning

We analysed *dmrad51* mutant adult flies for any phenotypic defect in the somatic tissues. Null mutant (*spnA093^A^*) as well as trans-heterozygote combinations showed defects in abdominal pattering, manifested as fusion of segments, segment polarity reversal and aberrant pigmentation of the cuticle (Fig. 2A2, marked with red arrow) as compared to the isogenic control (Fig. 2A1). Also, we observed consistent abnormalities in adult cuticle across various *dmrad51* alleles and trans-heterozygote combinations (Fig. 2A3-5, marked with red arrow). Knockdown of Dmrad51 with an RNAi specific for *dmrad51* driven by actin-Gal4 also recapitulated the adult cuticle phenotype (Fig. 2A6-A7, marked with red arrow). Further, we observed an inherent variability in the severity of the phenotype within a given genotype, which we categorised as extent of severity, ranging from S0 to S4 (where S0 corresponds to wild type like cuticle, whereas, S4 corresponds to the highest level of phenotypic severity) (Fig. 2B). We classified and quantified flies across *dmrad51* trans-allelic combinations for phenotypic severity. Within the same set of a given genotype, we observed a variable distribution of phenotypic severity (Fig. 2C), perhaps attributable to the degree of penetrance.

**Fig. 2.**
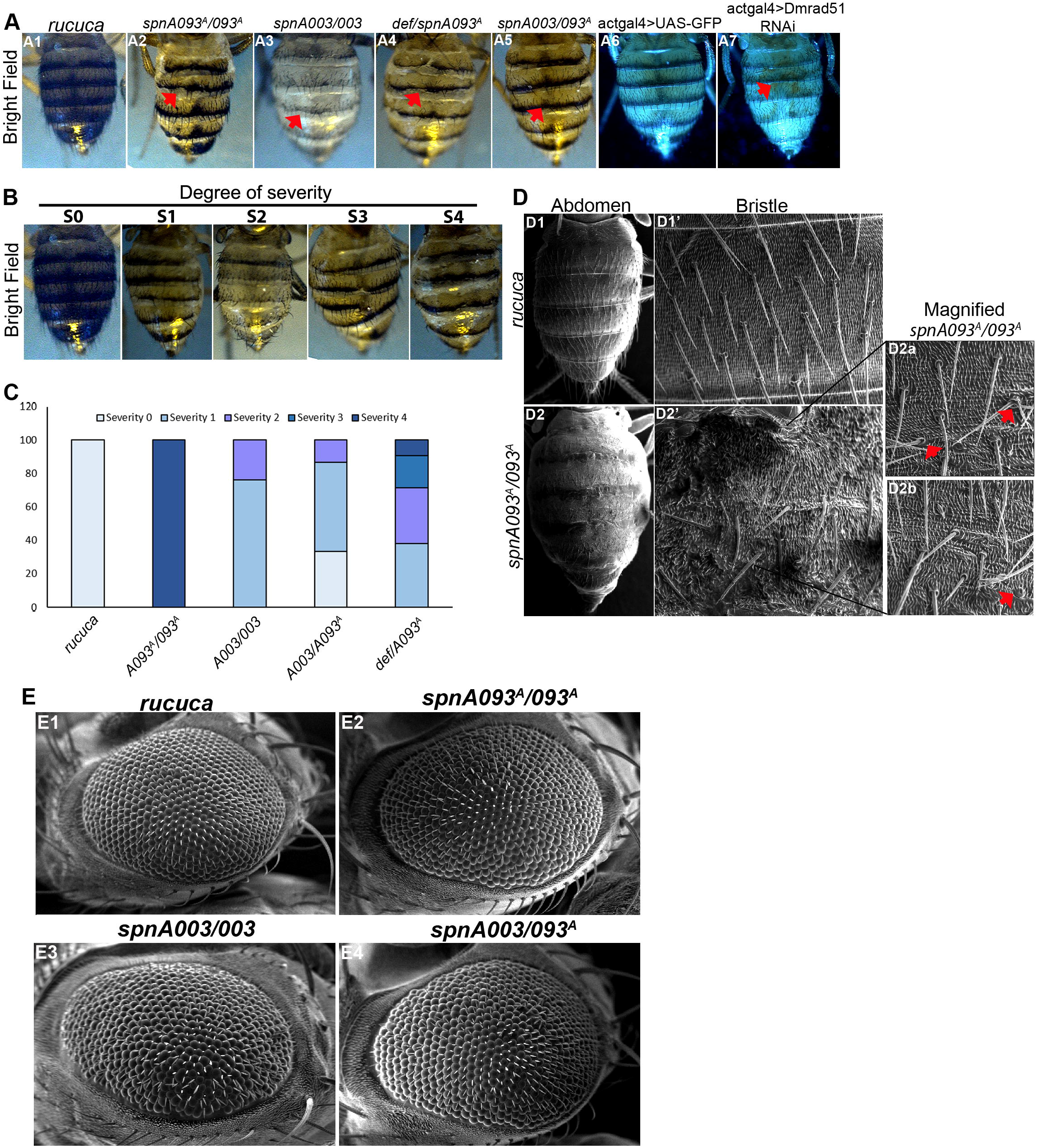
Absence of Dmrad51 function led to defects in adult cuticle patterning. (A1-A7) Bright field images of adult flies, control (A1), null mutant (*spnA093^A^*) (A2) and trans-heterozygotes (A3-A5), showing defects in abdominal stripes as evidenced by loss of pigmentation, and abnormal bristle patterns (red arrow marks the defect in the stripes). The same defect was recapitulated for Dmrad51 RNAi driven using act-Gal4 (A7), where Gal4 control is normal (A6). (B and C) Abdominal phenotype was graded on the basis of increasing severity of the defect from S0-S4 as shown in the bright field images (B) and the same is quantified across different genotypes (C). (D1-D2’) SEM analysis of abdominal cuticle showing patterned bristles in control (D1, high mag D1’), *dmrad51* mutant (D2, high mag D2’) exhibiting loss of bristle pattern and manifesting defects such as multiple bristles (D2a) and empty sockets (D2b). (E1-E4) SEM of adult eye control (E1), *dmrad51* null mutant (*spnA093^A^*) (E2) and trans-heterozygotes (E3-E4).

SEM analysis of the null mutant (*spnA093^A^*) abdominal cuticle showed defects in both macro- and microchaetae (Fig. 2D), evidenced by multiple bristles, loss of bristles polarity (Fig. 2D2a, marked with red arrow) and empty sockets (Fig. 2D2b, marked with red arrow). Our results indicate a possible dysregulation of the mechanisms involved in adult abdominal cuticle laydown in *dmrad51* mutant, a process which occurs by co-ordinated histoblast multiplication and larval epithelium replacement (Ninov et al. 2007). Moreover, similar phenotypic defects have already been reported for XPB (important player for transcription coupled repair) mutant (*hay*) (Fregoso et al. 2007), suggesting a general role of DNA damage repair proteins in patterning of tissues during somatic development as well, apart from their well described roles during *Drosophila* germline development.

In addition to the defects in adult cuticle, *dmrad51* null mutant (*spnA093^A^*) also exhibited a rough eye phenotype as shown by SEM analysis of the adult eye (Fig. 2E2), which was absent in isogenic control (Fig. 2E1). We also observed rough eye phenotype in other trans-allelic combinations (Fig. 2E3-4). We note that, in our study, we observed only subtle phenotypic defects in the adult wing of null mutant as well as in trans-allelic combinations (Fig. 3). Our results put together, suggest that Dmrad51 plays an important role in patterning of somatic tissue development, apart from its well-known function in egg patterning during meiosis. Moreover, the function of Dmrad51 appears to be more critical in abdomen cuticle patterning during pupal stages, whereas structures arising from larval precursors such as wing and eye show milder defects.

**Fig. 3.**
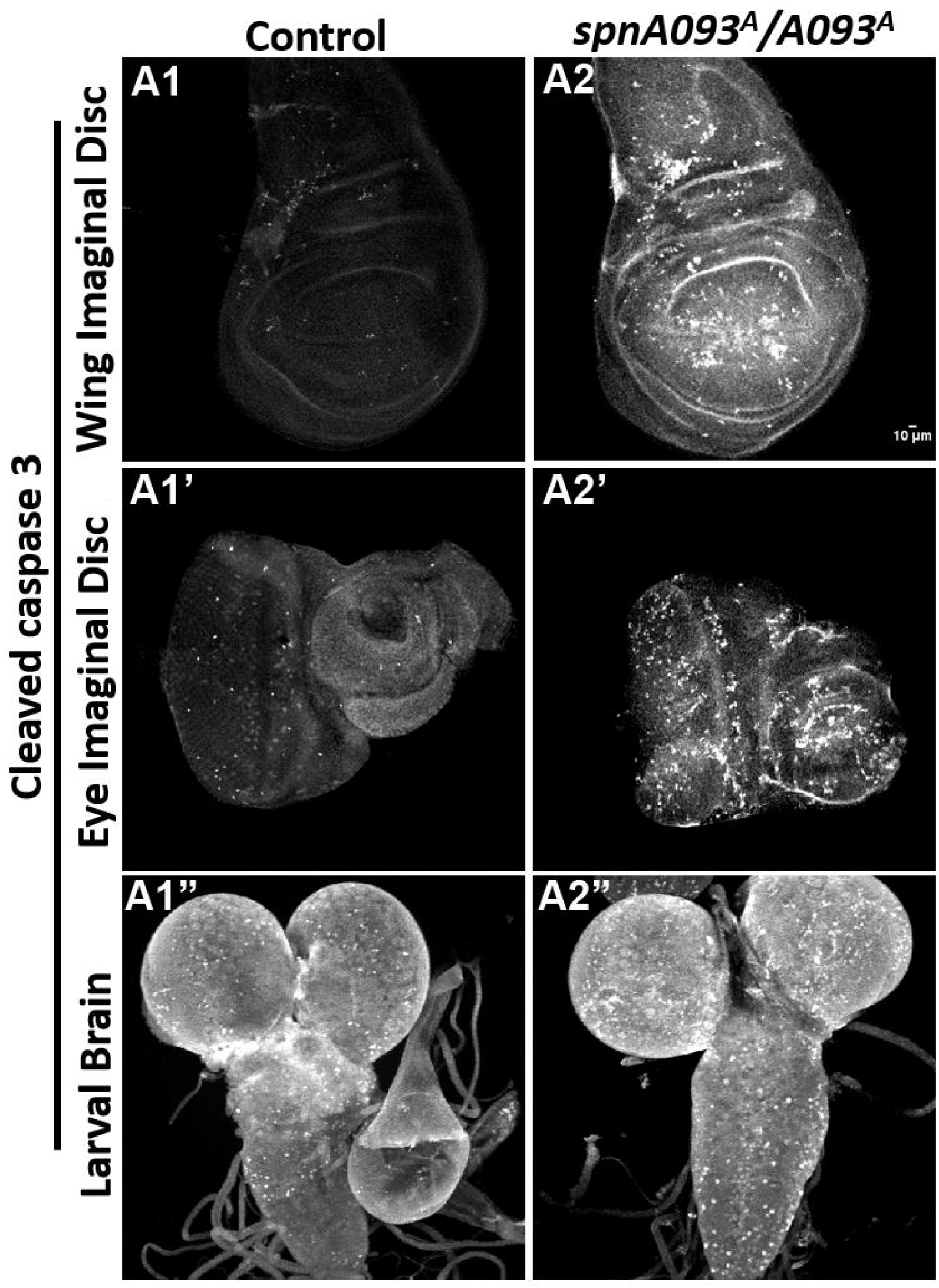
Larval structures show heightened level of cell death in *dmrad51 (spnA093^A^/093^A^*) mutant. (A1-A2) III instar wing discs assayed for CC3 in control wing disc (A1) and *dmrad51* null (A2). (A1’-A2’) III instar eye discs assayed for CC3 in control wing disc (A1’) and *dmrad51* null (A2’). (A1’’-A2’’) III instar brain assayed for CC3 in control wing disc (A1’’) and *dmrad51* null (A2’’).

### Wing imaginal disc and larval primordium exhibit increased levels of DNA damages and cell death but near normal adult wing phenotype

Since our analysis of mutant adult flies showed relatively milder phenotype in the structures arising from larval precursors such as eye and wing, we further probed for any signs of damages in their larval primordial structures. We showed that *dmrad51* null mutant (*spnA093^A^*) larval imaginal discs (wing as well as eye) (Fig. 3A2-A2’), showed an enhanced level of cell death as compared to the control (Fig. 31A1-A1’). A similar rise in cell death levels were observed in the mutant larval brain (Fig. 31A2”) in comparison to the control (Fig. 31A1”).

We further assayed levels of both cell death and DNA damages in mutant wing discs in different genetic combinations of *dmrad51* alleles. As expected, control wing disc exhibited normal wing morphology and there was only a basal level of cell death (Fig. 4A1 and B1-B1’). But despite a near normal morphology of the adult wing (Fig. 4A2), the null mutant (*spnA093^A^*) larval discs, showed a significant increase in cell death as assayed by cleaved caspase 3 (CC3) staining (Fig. 4B2), and quantified as percentage of CC3 positive areas (Fig. 4D). We also found our results to be consistent across trans-heterozygote groups of different *dmrad51* alleles, but with reduced intensity compared to the null mutant (Fig. 4A3-A5, B3-B5’ and D).

**Fig. 4.**
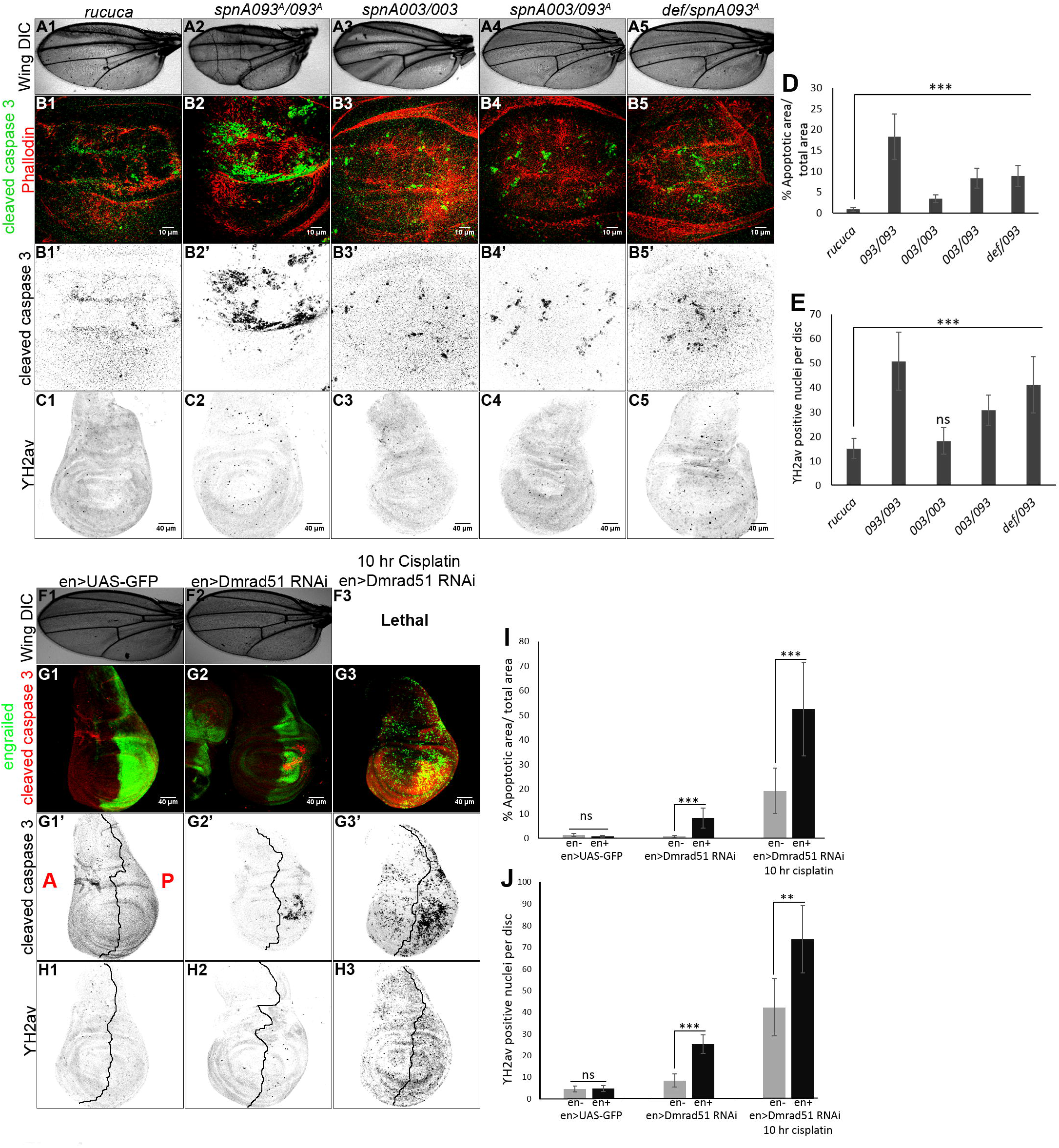
Wing imaginal disc and larval primordium exhibit increased levels of DNA damages and cell death but near normal wing phenotype. (A1-A5) DIC images of adult wing: control (A1), *dmrad51* null mutant (A2) and transheterozygotes (A3-A5). (B1-B5’ and D) III instar wing discs assayed for CC3 (green) and Phalloidin (red) (B1-B5); CC3 is also shown in grey scale (B1’-B5’). Control wing disc (B1-B1’), *dmrad51* null (B2-B2’) and trans-heterozygotes (B3-B5) (B3’-B5’). (D) Percentage of CC3 positive area quantitated by drawing a boundary manually around cleaved caspase 3 positive (high CC3) area using image J, and normalized to the total disc area as percentage (N= 15-20 wing disc from each genotype). (C) Wing discs assayed for YH2av, control (C1), *dmrad51* mutant (C2), trans-heterozygotes (C3-C5). (E) Quantification of the number of YH2av positive nuclei per disc (N= 15 wing disc from each genotype). We quantified only the intense-discrete YH2Av nuclei as YH2Av-positive nuclei and counted numbers using image J cell counter and expressed per whole disc. (F1-F3) DIC images of adult wing of Gal4 control (F1), Dmrad51 RNAi driven by en-Gal4 (F2). (G1-G3’) Wing discs assayed for CC3 (red) and en-Gal4 compartment (green), of Gal4 control UAS-GFP (G1), Dmrad51 RNAi (G2), and 10 hr cisplatin treated wing disc from Dmrad51 RNAi (G3) along with their corresponding grey scale images for CC3 (G1’-G3’). (I) Quantification of the percentage of CC3 positive areas (N= 12 wing disc from each genotype). (H1-H3) Wing discs assayed for YH2av belonging to Gal4 control (H1), Dmrad51 RNAi (H2), and 10 hr cisplatin treated wing disc from Dmrad51 RNAi (H3). (J) Quantification of the numbers of YH2av positive nuclei (N= 12 wing disc from each genotype). Data are mean ± SDM. *p < 0.05, **p < 0.01 and ***p < 0.001

In parallel, we assayed YH2av levels to monitor DNA damage response in the same system (Madigan et al. 2002; Khan et al. 2017). Null mutant wing discs showed higher YH2av positive nuclei (Fig. 4C2 and E) as compared to control (Fig. 4C1 and E). Trans-heterozygote allelic combinations also recapitulated a similar result (Fig. 4C3-C5 and E). We confirmed these results by RNAi knockdown using en-Gal4 driver which expresses in the posterior compartment of the wing discs. Expectedly, Dmrad51 knockdown showed significantly higher levels of cell death and YH2av in the posterior versus anterior compartment, without any effect on the adult wing (Fig. 4F2, G2-G2’, H2, I and J) as compared to Gal-4 control (Fig. 4F1, G1-G1’, H2, I and J). Moreover, compartments with Dmrad51 knockdown exhibited higher sensitivity to DNA damaging agent, cisplatin, compared to the internal control (Fig. 4F3, G3-G3’, H3, I and J). These results put together, suggest that Dmrad51 function is critical for cell survival in the somatic tissues, perhaps due to its important role in repairing endogenous DNA damages.

### *dmrad51* mutant cuticle shows low temperature sensitivity

Serendipitously, we observed an increase in severity of cuticle phenotype in *dmrad51* mutant at low temperature (18°C). This was in contrast to the fact that typically the mutant phenotypes tend to show heightened severity at higher temperature (29°C). This led us to perform a detailed analysis of *dmrad51* mutant at different temperatures, namely, 18°C, 25°C and 29°C. We saw a higher severity of abdominal cuticle phenotype at low temperature (18°C, Fig. 5A1-A1”) in comparison to high temperature (29°C, Fig. 5A2-A2”) across different *dmrad51* allelic trans-heterozygotes. More specifically, flies grown at 25°C (Fig. 5A3-A3”) exhibited milder severity than at 18°C, but more severe than at 29°C, (25°C phenotype was intermediate to 18°C and 29°C), thus showing an inverse correlation of phenotype with temperature increase. Further, we calculated the number of flies in different categories of severity ranging from S0 to S4 across various *dmrad51* trans-heterozygotes. Our analysis showed that 18°C led to an increase in the percentage of flies belonging to higher severity groups S4, S3 and S2, while the null mutant (*spnA093^A^*) was lethal at the same temperature (Fig. 5B). Interestingly at 29°C, we observed a higher percentage of flies in S0 and a small percentage of flies in S1 category across all the *dmrad51* alleles analysed including null mutant (*spnA093^A^*) (Fig. 5B). On the other hand at 25°C, we observed a relatively higher percentage of S1 but almost no S2, S3 and S4 categories in the transheterozygotes, while null mutant (*spnA093^A^*) mostly fell into the S4 category (Fig. 5B). Thus, our results taken together suggested that *dmrad51* mutant adult abdominal cuticle show a higher sensitivity at low temperature, conversely high temperature leads to the rescue of the phenotype.

**Fig. 5.**
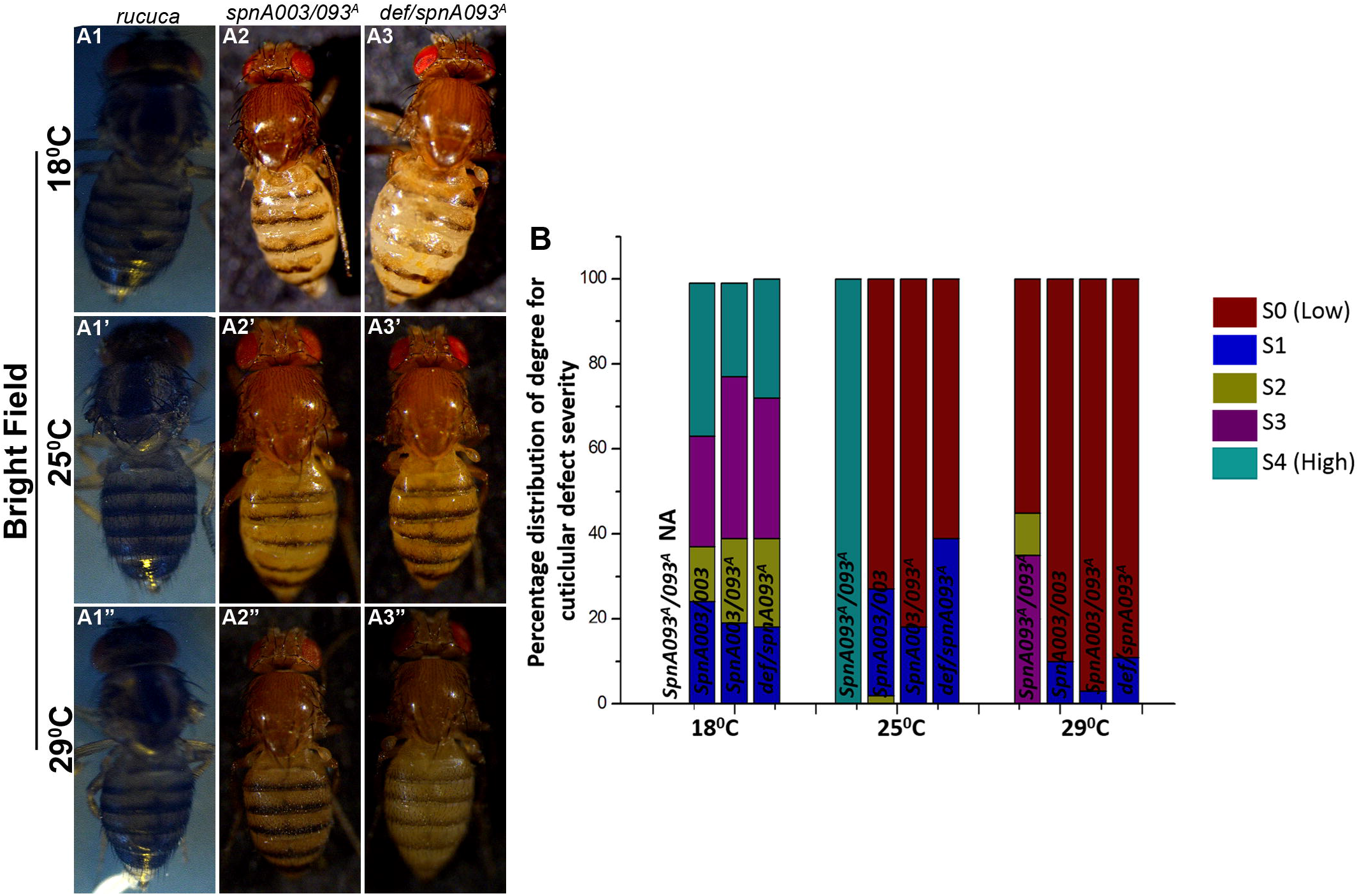
*dmrad51* mutant cuticle adult phenotype show low temperature sensitivity. (A1-A3”) Bright filed images of adult flies grown at different temperature, 18°C isogenic control (A1) trans-heterozygotes (A2-A3), 25°C isogenic control (A1’) transheterozygotes (A2’-A3’) and 29°C isogenic control (A1”) trans-heterozygotes (A2”-A3”). (B) Distribution of extent of severity ranging from (S0-S4, 18°C, 25°C and 29°C colour coded) at different temperature.

We also analysed structures arising from larval primordium (imaginal discs) adult wing and eye for temperature sensitivity. We performed an SEM analysis on adult eyes from flies grown at different temperatures (18°C, 25°C and 29°C); however unlike the abdominal cuticle, at low temperatures we did not observe an enhanced severity of the rough eye phenotype across various classes of the *dmrad51* trans-heterozygotes (data not shown). Also, we noted that the adult wings from *dmrad51* transheterozygotes did not show increased severity to low temperature (data not shown). Our results clearly revealed that the adult abdominal cuticle shows higher sensitivity to the absence of Dmrad51 at low temperature than the structures arising from larval primordium (wing and eye), thus suggesting a more critical role for Dmrad51 in adult cuticle patterning and that the process was sensitive to low temperature.

### Larval precursors of wing imaginal discs also exhibit heightened sensitivity to low temperature

Absence of Dmrad51 led to an increased sensitivity to low temperatures especially in the phenotype observed in abdominal cuticle, but not in the adult wing or eye. Therefore, we probed for signs of temperature sensitivity in larval precursors (III instar wing imaginal discs) of *dmrad51* mutant. Our analysis in *dmrad51* null mutant wing discs (*spnA093^A^*) as well as in trans-heterozygotes showed a significantly higher level of cell death at 18°C (Fig. 6A2-A5’) as compared to 25°C (Fig. 6C2-C5’, E and F), while controls showed no such increase in levels of cell death at low temperatures of 18°C (Fig. 6A1-A1’, C1-C1’, E and F). These results were in concordance with the low temperature sensitivity observed for adult cuticle phenotype and further validated that Dmrad51 absence is linked to low temperature sensitivity. However, the absence of major phenotypic defects in the adult wing even at 18°C can be accounted for by the robust compensatory pathways associated with wing imaginal discs as discussed later.

**Fig. 6.**
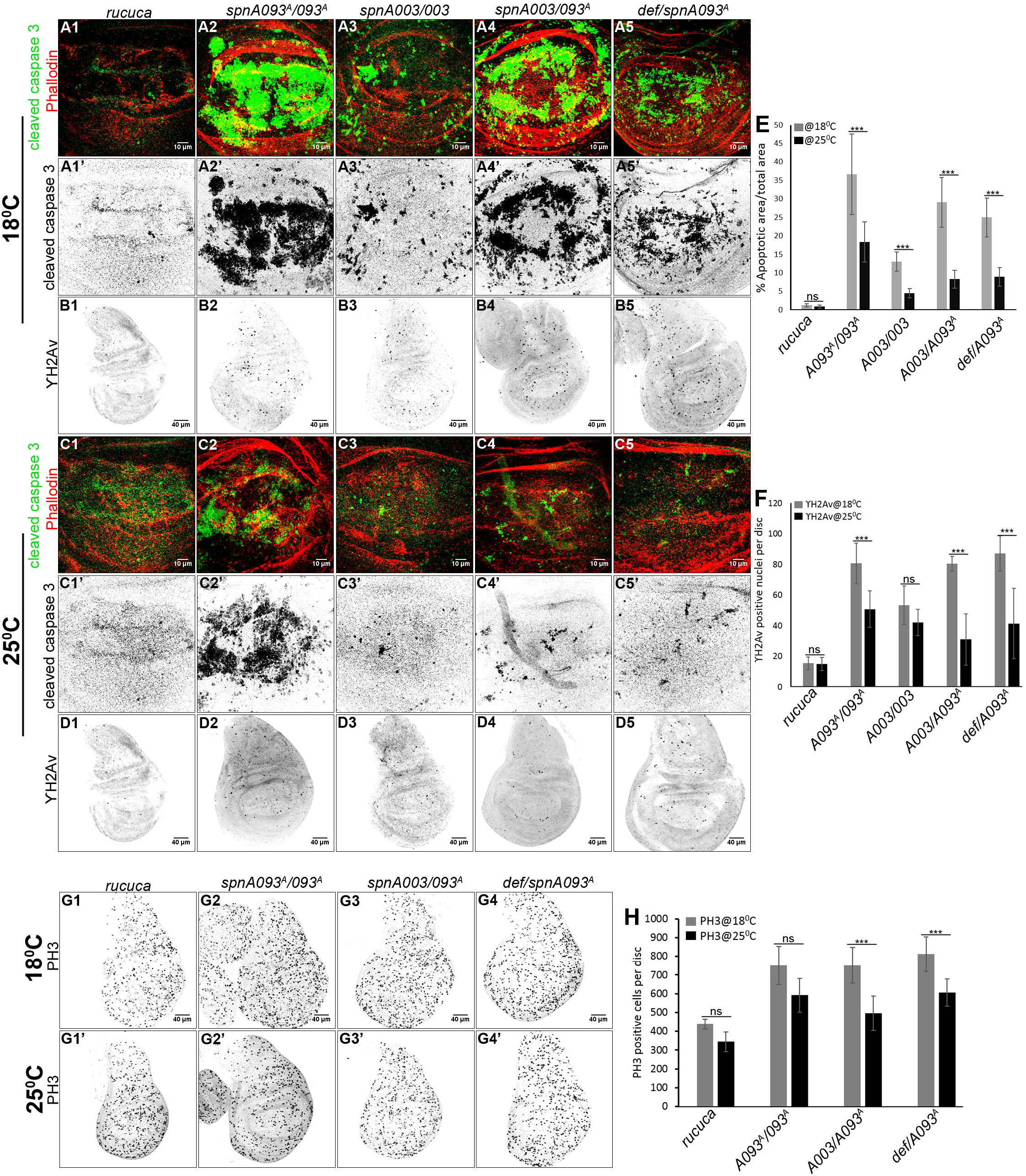
Larval precursor exhibits enhanced sensitivity to low temperature. (A1-A5’, C1-C5’ and E) III instar wing discs at 18°C (A1-A5’) and 25°C (C1-C5’) assayed for CC3 (green) and Phalloidin (red) (A1-A5 and C1-C5); CC3 is also shown in grey scale (A1’-A5’ and C1’-C5’). Control wing disc (A1-A1’ and C1-C1’), *dmrad51* null (A2-A2’ and A2-A2’) and trans-heterozygotes (A3-A5’, A3-A5’). Wing discs assayed for YH2av at 18°C (B1-B5) and 25°C (D1-D5), control (B1, D1), *dmrad51* mutant (B2, D2), trans-heterozygotes (B3-B5 and D3-D5). (E and F) Quantification of the percentage of CC3 positive areas and the number of YH2av positive nuclei per disc (N= 15 wing disc from each genotype). (G1-G4’ and H) III instar wing discs at 18°C (G1-G4) and 25°C (G1’-G4’) assayed for PH3 counts, which is quantitated as PH3 per disc (H). Data are mean ± SDM. *p < 0.05, **p < 0.01 and ***p < 0.001.

Further, we assayed for the levels of DNA damages in mutant wing imaginal disc using YH2av levels as readouts grown at 18°C and 25°C temperature. We observed increased levels of DNA damages in *dmrad51* mutant at 18°C (Fig. 6B2-B5 and F) as compared to 25°C (Fig. 6D2-D5 and F), whereas controls showed no such difference (Fig. 6B1, D1 and F). These results suggested that perhaps repair process is sensitive to the low temperature; however, this need to be studied carefully using other DNA repair mutants and also by exogenously induced DNA damages. In addition to increased cell death and DNA damages, we observed a higher level of mitotic counts in *dmrad51* mutant, as assayed for by phospho-Histone 3 (PH3) at 18°C compared to 25°C temperature. Further, these observations also point towards the possibility of abnormalities in mitosis or cell cycle at low temperatures and a possible role for Dmrad51 in cell cycle regulation, which we discuss below.

## DISCUSSION

In this study, we have investigated the role of Dmrad51 in somatic tissues during development, where absence of Dmrad51 leads only to subtle phenotypic defects in somatic tissues at adulthood, which was in sharp contrast to the rad51 function in higher organisms where its absence was associated with early zygotic lethality (Tsuzuki et al. 1996). Near normal phenotype of the *dmrad51* mutant was possibly due to efficient backup DNA repair pathways such as NHEJ, which can compensate for impaired HR in *dmrad51* mutant (Preston et al. 2006). In addition, rad51 loss of function in *dmrad51* mutant could also be compensated by other rad51 paralogs such as spn-B, spn-C and spn-D (Ghabrial et al. 1998; Abdu et al. 2003; Radford & Sekelsky 2004). We report that *dmrad51* mutant show higher levels of cell death and damages during the larval stages, whereas adult structures show near normal morphology, perhaps reflecting high tissue regenerative capacity (Haynie & Bryant 1977) in *dmrad51* mutant. A similar phenotypic compensation has been shown in *Drosophila* centrosomal mutant, where damages were masked by cell death induced JNK activation (Poulton et al. 2014). Some of our unpublished results on wing imaginal discs also indicate a critical feedback regulatory role of dp53 and JNK pathway in cell death versus cell protection functions, respectively, which could account for the phenotypic compensation in adult wing, whose detailed mechanistic crosstalk is being explored currently.

We also performed lethality analysis using *spnA093^A^/def* trans-heterozygote combination, previously used to uncover germline defects (Staeva-Vieira et al. 2003). To our surprise, we did not observe any significant lethality in *spnA093^A^/def,* suggesting a possibility (data not shown), that in the *def* line absence of other genes (background of the stock) leads to a rescue of lethality caused by *dmrad51* mutant or that the *def* line only partially uncovered *dmrad51* mutant. Moreover, *spnA093^A^/def* failed to uncover rough eye phenotype observed in case of null mutant as well as other trans-heterozygote combinations (data not shown). In support of this, we observed that *spnA093^A^/def* female flies showed significant fertility as opposed to complete infertility in other *dmrad51* mutant alleles and allelic combinations (data not shown), suggesting that *def* only partially uncovers *dmrad51* mutant phenotypes.

There are very few instances in the literature which describe how low temperature can enhance the phenotypic severity; such rare mutants being referred to as cold sensitive mutants (cs-mutants) (Awano et al. 2007; Strauss & Guthrie 1991), whereas there are multiple instances of high temperature sensitivity causing mutations (ts-mutants), both being typically linked to temperature sensitive changes in protein structure at restrictive temperatures (Baliga et al. 2016). Interestingly, low temperature sensitivity observed in *dmrad51* mutant arises not due to any protein structural change, but rather due to the absence of the protein, Rad51 (functionally null because of premature termination of the protein), *per se*. We speculate that temperature sensitivity in *dmrad51* mutant could possibly arise due to some important cellular process(es) requiring Dmrad51 function which happens to be temperature sensitive. This could possibly arise due to sensitivity of DNA damage repair to low temperature, as mutant show higher levels of DNA damages at 18°C. Moreover, temperature sensitivity can also arise during cellular proliferation via the cell cycle checkpoint role of Dmrad51, as described for yeast rad51 (Symington 2002). Several DNA repair proteins have been implicated in centrosome stabilization in concert with their nuclear integrity function (Cappelli et al. 2011), absence of which might contribute to mitotic defects in dmrad51 mutant, which in turn could be temperature sensitive. All of possibilities deserve a detailed investigation to uncover the process sensitive to low temperature in the absence of Dmrad51. We suggest that the *dmrad51* mutant might provide an interesting system to probe mechanistically such temperature sensitive cellular processes, not associated with changes in the protein structure, an area of cell biology that is relatively very nascent now. Finally, we conclude that our current modular and descriptive report provides a viable and potentially interesting model system to uncover novel cellular functions of rad51, beyond its well prescribed roles in genomic recombination, using a genetically tractable organism, *Drosophila*.

## MATERIAL AND METHODS

### Fly Stocks

Details are listed in Fig. 1B.

### Lethality Assay

Embryos from heterozygote cross [balanced with a GFP balancer (discussed in fly genetics)], were aligned after 6 hours of egg laying on sucrose agar plates in Halocarbon oil (HC-700-Sigma H8898). Mutant embryos were sorted on the basis of absence of GFP signal and scored for the hatching (empty chorion) or absence of hatching (dead embryo) after 36 hours. Percentage embryonic lethality was calculated by dividing total dead embryos to number of mutant embryo. For larval and pupal lethality, first instar larvae were transferred to cut-vials (containing fly media ~5ml) from sucrose agar plate, and then kept at 25°C for growing. Percentage of larval lethality was calculated as: Total number of larvae transferred (n=40) – Number of pupae/ Total number of larvae transferred (n=40). Similarly, pupal lethality was computed as: Total number of pupae – adult flies eclosed/ Total number of pupae.

### Adult Fly Imaging and Scanning Electron Microscopy

Adult flies were grown at respective temperatures (18°C, 25°C and 29°C) and aged for 4-5 days before imaging. In all the experiments, phenotype was scored only in female flies. Adult wing was clipped from 4-5 day old adult fly and DIC imaging was performed using Olympus FV1000. Adult cuticle, from the clipped wing, was photographed using Leica-Epifluorescence microscope (M165 FC) and Leica camera (DFC 7000 T). SEM analysis of adult eye was performed using Zeiss Scanning electron microscope at 500X magnification and 5.0KV beam.

### Immunohistochemistry and Imaging

Imaginal discs were fixed using 4% formaldehyde in 1XPBS, followed by blocking with 0.5% BSA dissolved in 0.1% PBST for 2 hr. Primary antibody incubations were done at 4°C overnight, followed by incubation with secondary-Ab for 2 hr at room temperature. Following antibody incubation, all washes were done in 0.1% PBST [Primary antibodies: CC3 (Sigma-8487, 1:200), active-PH3 (Santacruze-9578, 1:500), YH2Av (DSHB, 1:50), engrailed (DSHB, 1:50), anti GFP (DSHB, 1:50)]. Alexa fluor secondary antibodies were used at 1:200 dilutions. Image acquisition was done on Olympus FV 1000, intensity adjustments were done using Photoshop PS5 and Image J was used for image processing.

## ACKNOWLEDGMENTS

We acknowledge Trudi Schupbach (Princeton University) for Dmrad51 stocks; Ms. B Chalke and R D Bapat from TIFR-SEM facility; Profs. Maithreyi Narasimha and Ullas Kolthur from TIFR for critical inputs; Bloomington Stock Center, Developmental Studies Hybridoma Bank, and Vienna *Drosophila* Resource Center for reagents; and Funding support to BJR from DAE-TIFR and JC Bose award grant (DST).

## References

Abdu U. et al., 2003. The Drosophila spn-D Gene Encodes a RAD51C-Like Protein That Is Required Exclusively During Meiosis.

Awano N. et al., 2007. Complementation Analysis of the Cold-Sensitive Phenotype of the Escherichia coli csdA Deletion Strain □., 189(16), pp.5808–5815.

Baliga C. et al., 2016. Rational elicitation of cold-sensitive phenotypes.

Baumann, P. & West, S.C., 1998. Role of the human RAD51 protein in homologous recombination and double-stranded-break repair. Trends in Biochemical Sciences, 23(7), pp.247–251. Available at: http://www.sciencedirect.com/science/article/pii/S0968000498012328.

Cappelli E. et al., 2011. Homologous recombination proteins are associated with centrosomes and are required for mitotic stability. Experimental Cell Research, 317(8), pp.1203–1213. Available at: http://dx.doi.org/10.1016/j.yexcr.2011.01.021.

Ghabrial, A., Ray, R.P. & Schüpbach, T., 1998. okra and spindle-B encode components of the RAD52 DNA repair pathway and affect meiosis and patterning in Drosophila oogenesis. Genes and Development, 12(17), pp.2711–2723.

Haynie, J.L. & Bryant, P.J., 1977. The effects of X-rays on the proliferation dynamics of cells in the imaginal wing disc of Drosophila melanogaster. Wilhelm Roux’s Archives of Developmental Biology, 183(2), pp.85–100.

Helleday, T., Eshtad, S. & Nik-zainal, S., 2014. Mechanisms underlying mutational. Nature Publishing Group, 15(9), pp.585–598. Available at: http://dx.doi.org/10.1038/nrg3729.

Jeggo, P.A. & Lo, M., 2007. DNA double-strand breaks: their cellular and clinical impact?, pp.7717–7719.

Kelso A.A. et al., 2017. Data in Brief Data on Rad51 amino acid sequences from higher and lower eukaryotic model organisms and parasites. Data in Brief, 10, pp.364–368. Available at: http://dx.doi.org/10.1016/j.dib.2016.12.002.

Khan C. et al., 2017. DNA damage signalling in D. melanogaster requires non-apoptotic function of initiator caspase Dronc., (July).

Madigan, J.P., Chotkowski, H.L. & Glaser, R.L., 2002. DNA double-strand break-induced phosphorylation of Drosophila histone variant H2Av helps prevent radiation-induced apoptosis. Nucleic acids research, 30(17), pp.3698–3705.

Mao Z. et al., 2008. Brief report Comparison of nonhomologous end joining and homologous recombination in human cells., 7, pp.1765–1771.

Mao Z. et al., 2017. DNA repair by nonhomologous end joining and homologous recombination during cell cycle in human cells ND ES SC RIB., 4101 (July).

Petrini, J.H., Bressan, D.A. & Yao, M.S., 1997. The RAD52 epistasis group in mammalian double strand break repair. Seminars in immunology, 9(3), pp. 181 – 8. Available at: http://www.sciencedirect.com/science/article/pii/S1044532397900671.

Poulton, J.S., Cuningham, J.C. & Peifer, M., 2014. Acentrosomal Drosophila epithelial cells exhibit abnormal cell division, leading to cell death and compensatory proliferation. Developmental cell, 30(6), pp.731–45. Available at: http://www.ncbi.nlm.nih.gov/pubmed/25241934.

Preston, C.R., Flores, C.C. & Engels, W.R., 2006. Differential usage of alternative pathways of double-strand break repair in Drosophila. Genetics, 172(2), pp.1055–1068.

Radford, S.J. & Sekelsky, J.J., 2004. Taking Drosophila Rad51 for a SPiN. Nature structural & molecular biology, 11 (January), pp.9–10.

Shinohara, A., 1992. Rad51 Protein Involved in Repair and Recombination in S. cerevisiae Is a RecA-like Protein., 69, pp.457–470.

Staeva-Vieira, E., Yoo, S. & Lehmann, R., 2003. An essential role of DmRad51/SpnA in DNA repair and meiotic checkpoint control. EMBO Journal, 22(21), pp.5863–5874.

Staeva-Vieira, E., Yoo, S. & Lehmann, R., 2003. An essential role of DmRad51 / SpnA in DNA repair and meiotic checkpoint control., 22(21), pp.5863–5874.

Strauss, E.J. & Guthrie, C., 1991. A cold-sensitive mRNA splicing mutant is a member of the RNA helicase gene family., pp.629–641.

Symington, L.S., 2002. Role of RAD52 Epistasis Group Genes in Homologous Recombination and Double-Strand Break Repair., 66(4), pp.630–670.

Tearle, R.G. & Nusslein-Volhard, C., 1987. Tubingen mutants and stock list. Drosophila Information Service, 66, pp.209–269.

Translocations H. et al., 2001. Saccharomyces cerevisiae rad51 Mutants Are Defective in DNA Damage-Associated Sister Chromatid Exchanges but Exhibit Increased Rates of., 1.

Tsuzuki T. et al., 1996. Targeted disruption of the Rad51 gene leads to lethality in embryonic mice. Proceedings of the National Academy of Sciences of the United States of America, 93 (June), pp.6236–6240.

Yoo, S., 2006. Characterization of Drosophila Rad51/SpnA protein in DNA binding and embryonic development. Biochemical and Biophysical Research Communications, 348(4), pp. 1310–1318.

Yoo, S. & McKee, B.D., 2005. Functional analysis of the Drosophila Rad51 gene (spn-A) in repair of DNA damage and meiotic chromosome segregation. DNA Repair, 4(2), pp.231–242.

